# Insights from *Melipona bicolor* hybrid genome assembly: A stingless bee genome with chromosome-level scaffold

**DOI:** 10.1101/2023.10.09.561533

**Authors:** Natalia de Souza Araujo, Fernando Ogihara, Pedro Mariano Martins, Maria Cristina Arias

## Abstract

**Background:** The highly eusocial stingless bees are crucial pollinators of native and agricultural ecosystems. Nevertheless, genomic studies within this bee tribe remain scarce. We present the genome assembly of the stingless bee *Melipona bicolo*r. This bee is a remarkable exception to the typical single-queen colony structure, since in this species, multiple queens may coexist and share reproductive duties, resulting in genetically diverse colonies with weak kinship connections. As the only known genuinely polygynous bee, *M. bicolor*’s genome provides a valuable resource for investigating sociality beyond kin selection.

**Results:** The genome was assembled employing a hybrid approach combining short and long reads, resulting in 241 contigs spanning 259 Mb (N50 of 6.2 Mb and 97.5% complete BUSCOs). Comparative analyses shed light on some evolutionary aspects of stingless bee genomics, including multiple chromosomal rearrangements in *Melipona*. Additionally, we explored the evolution of venom genes in *M. bicolor* and other stingless bees, revealing that, apart from two genes, the conserved repertoire of venom components remains under purifying selection in this clade.

**Conclusion:** This study advances our understanding of stingless bee genomics, contributing to the conservation efforts of these vital pollinators and offering insights into the evolutionary mechanisms driving their unique adaptations.

## Background

The Meliponini, commonly known as the stingless bees, comprise a diverse group of highly eusocial bees that play a crucial role as pollinators of native and crop plants [1–3]. These bees are essential for the sustainable development of local ecosystems, particularly in the Neotropics, as they contribute to the maintenance of native flora diversity while also enhancing agricultural productivity [3, 4]. Nonetheless, the diversity of stingless bees faces constant threats for multiple reasons such as habitat loss, inappropriate beekeeping practices, and the introduction of invasive species[1, 5–7]. Stingless bees also serve as valuable models for behavioral and evolutionary studies given their wide range of adaptations [8] as the tribe comprises about 500 species and 48–61 genera [9, 10]. Despite the importance of the group, genomic studies in Meliponini are scarce and only nine stingless bee genomes have been made available on NCBI [11], none of which were based on long-read sequencing. In an effort to diminish this knowledge gap, we performed the genome sequencing, assembly, and annotation of the stingless bee *Melipona bicolor* (Lepeletier, 1836), using a hybrid assembly approach that combined short and long reads data. To gain some insights into stingless bees’ genomic evolution, we comparatively analyzed this genome with other corbiculate bee species.

The nuclear genome of *M. bicolor* has the typical chromosome number for *Melipona* species (n = 9) [13] and an estimated genome size of 273.84 Mb [14]. Moreover, *M. bicolor* is part of the so-called Group I of *Melipona* species characterized by low heterochromatin content [13, 14]. Alike honeybees, almost all stingless bee species exhibit monogynic perennial colonies, where a single reproductive queen resides alongside multiple workers [9]. An intriguing exception to this rule is observed in *Melipona bicolor* [15], an endemic bee species found in the Brazilian Atlantic Rain Forest [10]. In *M. bicolor* colonies, multiple queens may co-exist and share the reproductive duties of the colony [15, 16]. Notably, polygyny is not mandatory in *M. bicolor*, and the number of queens within a colony can fluctuate [16]. Additionally, workers in *M. bicolor* colonies may accept queens from different genetic backgrounds [17], and they may contribute to male production by laying unfertilized eggs [18]. As a result, polygynic colonies of *M. bicolor* exhibit greater genetic diversity and weak kinship connections [17, 19]. This unique characteristic of *M. bicolor* makes it an exceptionally interesting model for investigating the maintenance of sociality beyond kin selection. Future studies based on its genome will be able to further delve into these aspects, providing valuable insights into the complex dynamics of social behavior in this species.

One of the distinguishing features of stingless bees, as their name suggests, is their inability to sting and inject venom because their stinging apparatus is stunted [20]. The sting apparatus is a modified ovipositor found in Hymenoptera, that allows the inoculation of venom into prey and/ or aggressors [21, 22]. The bee venom of the Apini and the Bombini have been extensively characterized, revealing the presence of various substances such as melittin, apamin, hyaluronidase, phospholipase A2, and venom allergens [23–27]. Even though the composition of most venoms is known to be complex and to rapidly vary according to diverse factors such as age, diet, and sex [28, 29], in a comparative study across the Hymenoptera, Koludarov et al. (2022) found that most venom genes are conserved throughout the group. Intriguingly, these authors also reported that, except for the *melittin* gene, stingless bees still have a complete gene repertoire of bee venom components in their genomes. Herein, we incorporate the genome of *M. bicolor* to further investigate the evolutionary trajectory of venom-associated genes within stingless bees.

By expanding our understanding of the genomic landscape of Meliponini, this study contributes to the conservation and sustainable management of these important pollinators. Additionally, it will enable future studies aiming at understanding the evolutionary processes that have shaped the diverse traits and adaptations observed within this important bee group.

## Results

### *M. bicolor* genome assembly

We generated 624,925,618 short and paired reads of 100 bp in length and 1,628,680 long subreads with N50 of 13,649 bp. The Falcon assembler, based on long reads only, resulted in an assembly of 249,831,102 bp total size with an N50 of 466,343 bp across 1,048 contigs, while the hybrid assembly with MaSuRCA generated a genome with 277 contigs, of N50 5,245,763 bp and total length of 261,548,481 bp. After merging these two assemblies, scaffolding with transcriptomic data, and polishing we obtained an assembly with 259 contigs, N50 of 6,225,681 bp, and a total length of 260,232,714 bp. From this, we removed one mitochondrial contig and 17 other contigs with low support (i.e., <1x of long read coverage) resulting in our final genome assembly with 241 contigs totalizing 259,858,556 bp with an N50 of 6,225,681 bp, L50 of 12, and largest contig size of 36,767,525 bp (Table SI). This final genome assembly and its annotation, as well as all datasets used to generate them, are available at NCBI under the BioProject PRJNA632864, and accession code JAHYIQ01.

The mean coverage of long and short reads was respectively 42 (±15 SD) and 234 (±626 SD). Sequencing coverage was congruent between the two sequencing strategies (Figures S1 and S2). No contaminants were identified through the BlobToolKit analyses as all identified matches to databases were against other bee species (Figure S3). Nevertheless, almost all smaller contigs have a distinct GC proportion when compared to the larger contigs. Small contigs were mostly GC-rich (Figure 1, S3), which may indicate that these contigs are most likely composed of repetitive DNA. As seen in Figure 1, the largest scaffold reached a chromosome size of over 36 Mb for which the long reads were essential to assembly. Notably, keeping contiguity in low GC areas (Figure S1).

**Figure 1.**
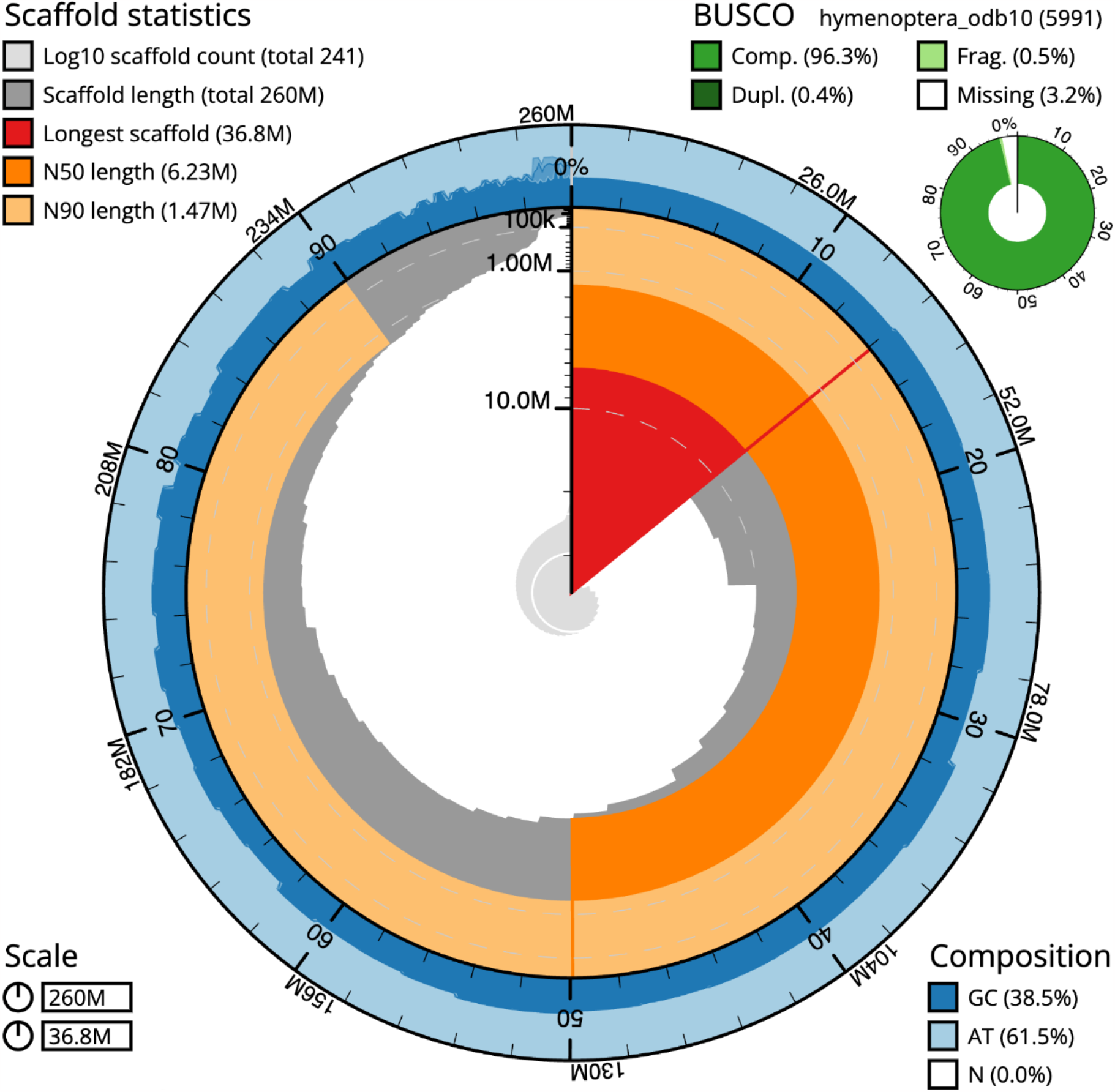
Summary representation of *M. bicolor* genome assembly. The main snailplot is divided into 1,000 size-ordered bins, each representing 0.1% of the total assembly size (259,858,556 bp). Sequence length distribution is shown in dark grey scaled by the largest scaffold of 36,767,525 bp, shown in red). Orange and pale orange indicate the N50 (6,225,681 bp) and N90 (1,474,512 bp), respectively. The pale grey spiral shows the cumulative sequence count on a log scale with white scale lines showing successive orders of magnitude. The blue and pale blue area shows GC and AT distributions. A summary of complete, fragmented, duplicated, and missing BUSCO genes in the hymenoptera_odb10 set is shown in the top right.

Against the hymenoptera_odb10 (5,991 BUSCO orthologs) the genome of *M. bicolor* contained 96.3% complete BUSCOs (single: 95.9%, duplicated: 0.4%), and 0.5% fragmented and 3.2% missing BUSCO orthologs, respectively. These results are comparable to the best bee genomes available (Table I), including that of the honeybee and the bumblebee. In terms of contiguity, *M. bicolor* assembly represents a significant improvement from other stingless genome assemblies available with a much larger N50 and fewer contigs (Figure 2).

**Table 1.** BUSCO comparative analyses of conserved orthology across two high-quality genome assemblies (*B. terrestris* and *A. mellifera*), *M. bicolor* assembly (in bold), and other better-quality stingless bee genomes available at NCBI.

| Species | Complete (%) | Single (%) | Duplicated (%) | Fragmented (%) | Missing (%) |
| --- | --- | --- | --- | --- | --- |
| *Apis mellifera* | 97.7 | 97.6 | 0.1 | 0.3 | 2 |
| *Bombus terrestris* | 96.5 | 96.3 | 0.2 | 1.2 | 2.3 |
| *Melipona quadrifasciata* | 96.5 | 96.3 | 0.2 | 1.2 | 2.3 |
| *Frieseomelitta varia* | 96.4 | 96.2 | 0.2 | 1.2 | 2.4 |
| ***Melipona bicolor*** | **96.3** | **95.9** | **0.4** | **0.5** | **3.2** |
| *Tetragonula mellipes* | 88.8 | 88.5 | 0.3 | 5.5 | 5.7 |
| *Tetragonula davenporti* | 87.9 | 87.6 | 0.3 | 6.3 | 5.8 |
| *Tetragonula carbonaria* | 86.9 | 86.6 | 0.3 | 6.7 | 6.4 |
| *Tetragonula clypearis* | 85.7 | 85.4 | 0.3 | 7.5 | 6.8 |
| *Tetragonula hockingsi* | 85.4 | 85.1 | 0.3 | 7.3 | 7.3 |

**Figure 2.**
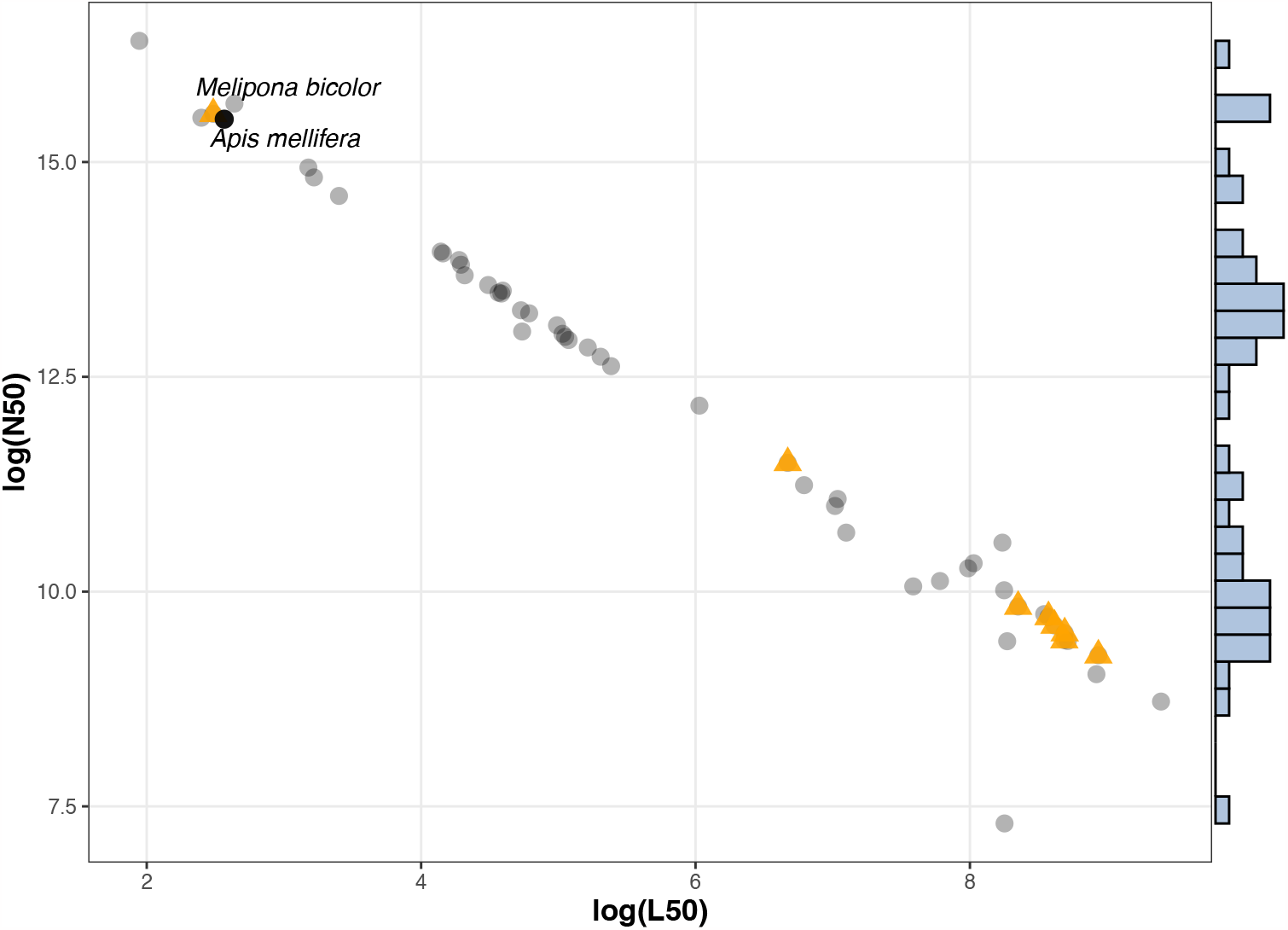
N50 and L50 values of all bee genomes available at NCBI in June 2021. Values are shown in the log base. Stingless bee genomes are indicated by orange triangles, the honeybee genome is represented by a black-filled circle, while all other bee species (49 genomes) are represented by grey circles. Cumulative frequencies are shown in the steel blue bars.

### *M. bicolor* genome annotation

Repetitive regions accounted for 18% of the genome. Most repeats were unclassified (11.5%) or non-interspersed (4%), while transposable elements of Class 1 and Class 2 corresponded to 1.5% and 1.6%, respectively. The complete classification of repetitive elements observed in the genome follows in Figure 3. The repetitive elements final library and report are available at (https://github.com/nat2bee/repetitive_elements_pipeline/tree/master/database). After repeating masking, 20,428 gene models were identified in the genome resulting in the annotation of 21,371 protein-coding genes and 150 tRNAs. Functional annotations were obtained for 15,010 of these protein-coding genes using the Funnannotate pipeline, while 11,905 *M. bicolor* genes were blasted against the bee genes in the UniRef90 database.

**Figure 3.**
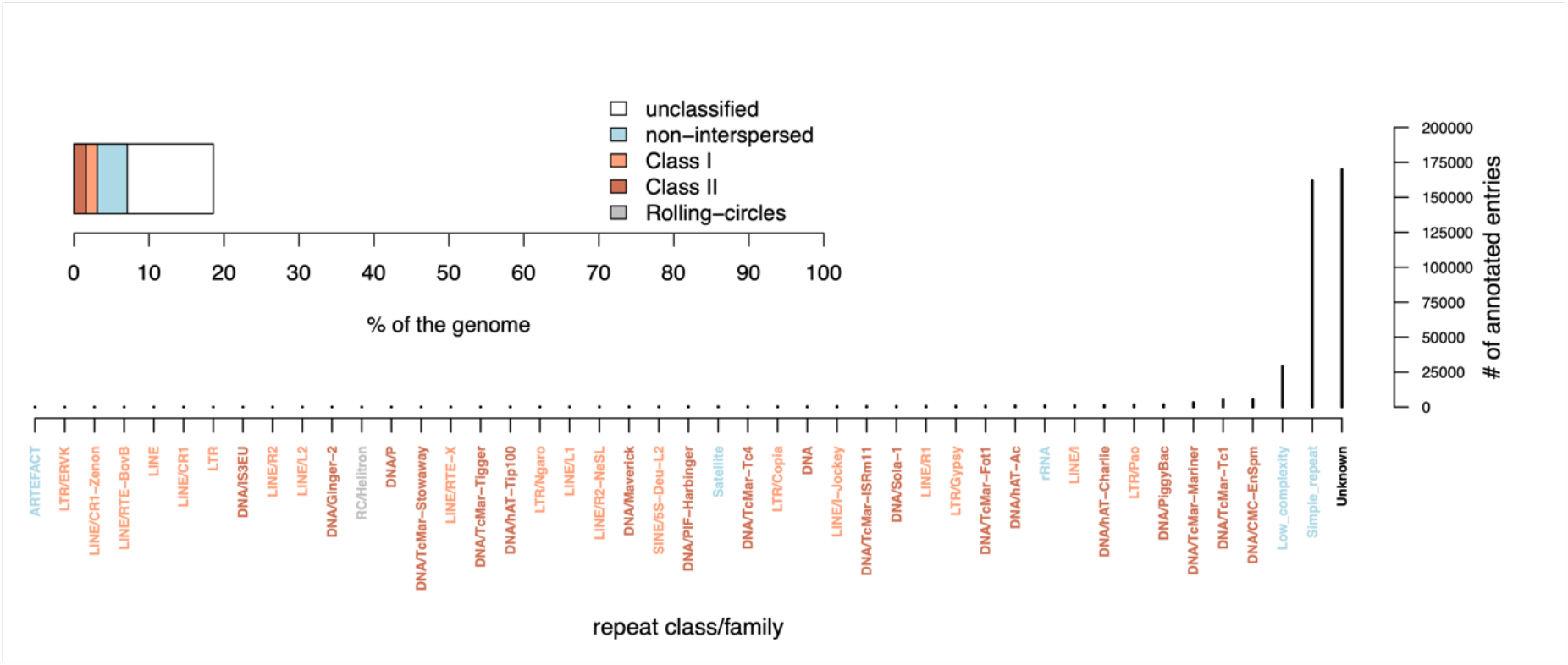
Repetitive elements found in the genome of *M. bicolor*. Repetitive regions accounted for about 18% of the genome assembly (upper plot). The lower plot shows the frequencies of all identified repeats with more frequent categories closer to the y-axis.

### Comparative genomics

Among the twelve corbiculate bees (Table SI) and the outgroup species *H. laboriosa*, we found 12,514 orthogroups clustering 164,535 genes (91.4% of the total). The largest 4,440 orthogroups contained 14 or more genes, and 6,760 orthogroups included representative genes from all species, with 2,826 of these consisting of orthogroups of single-copy genes (all genes contained in orthogroups follow in Supplementary File 2). Although *M. bicolor* had the largest number of analyzed proteins, only 64.9% (or 12,777) of them were assigned to orthogroups (Figure 4). Still, 87% of the orthogroups included *M. bicolor* genes and the proportion of species-specific genes for this species was not as high as that of other stingless bees (0.9%). Nineteen orthogroups were exclusive to eusocial corbiculate species and occurred in *M. bicolor* (Table SII), but not necessarily in other stingless bee genomes (15 also occurred in *M. quadrifasciata*, and 8 in *F. varia*). These orthogroups were associated with biological functions related to *nucleosome assembly, phospholipid metabolic process, lipid catabolic process, defense response to bacterium*, and *arachidonic acid secretion*.

**Figure 4.**
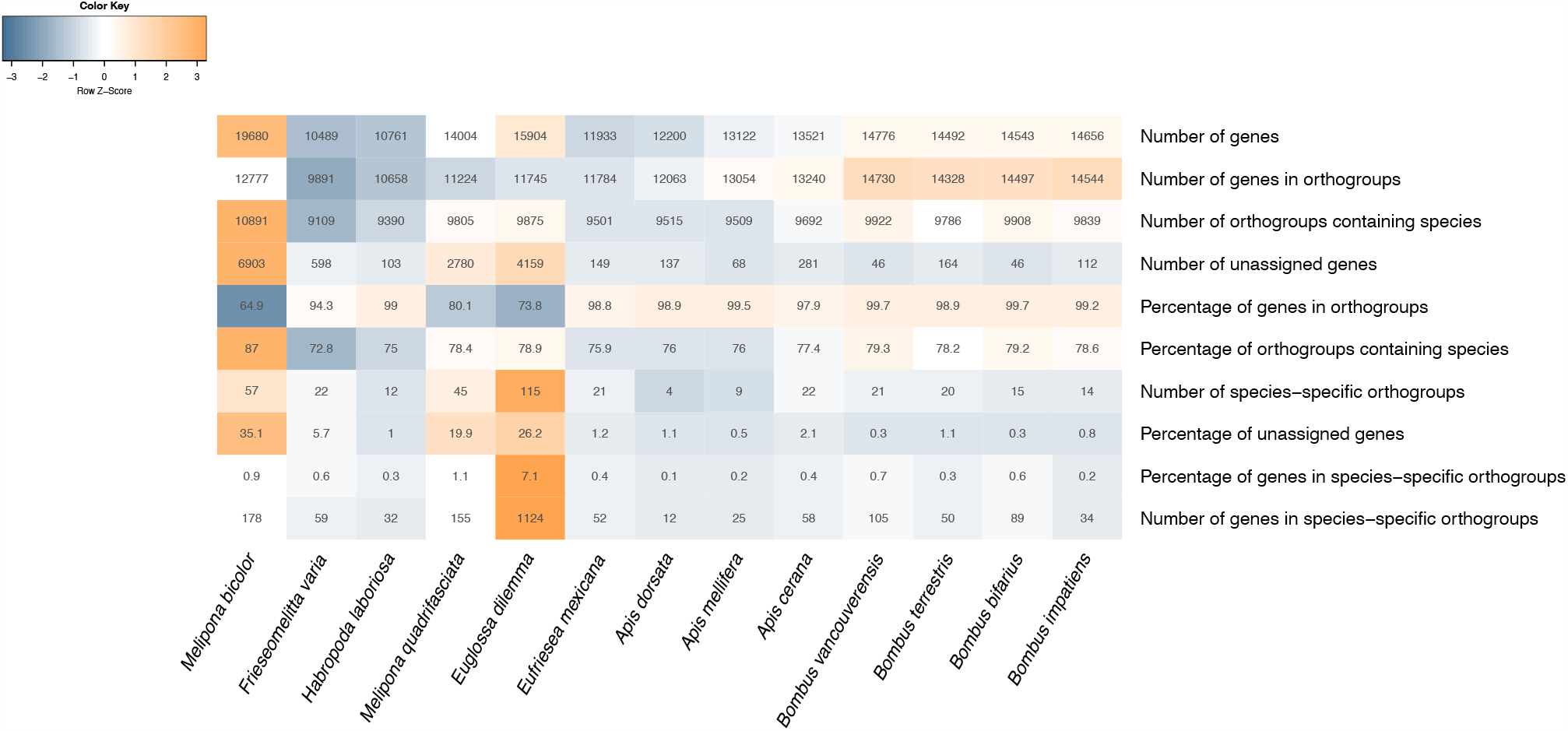
Summary results from OrthoFinder orthology analyses. Darker orange cells indicate larger values while darker shades of blue show smaller values compared to the others shown in the table.

Based on the 6,760 orthogroups including all species the estimated ultrametric species tree rooted at *H. laboriosa* had an average distance from root to leaves of 0.141434 (Supplementary File 1). Based on this tree and 12,548 orthologous genes (after excluding genes from large gene families) changes in gene family numbers (expansion or contraction) were identified. We found 148 gene families under significant changes across the studied species (p-value < 0.05, Supplementary File 3). Notably, in the nodes leading to the bumblebees and the honeybees, the great majority of changes led to increases in gene family sizes (Figure 5). In the node leading to all eusocial corbiculate, five gene families associated with *carbohydrate metabolic process, lipid metabolic process, fatty acid biosynthetic process*, and *signal transduction* were found to be in expansion, while only one unknown gene family was reported to have contracted, and it did not have an associated function (Table SIII). These results highlight the relevance of alterations in metabolism and immunity in the evolution of eusocial corbiculate bees. The branch leading to *M. bicolor* had 12 gene families significantly changing (10 increasing and 2 decreasing, Figure 5), these changes involved genes associated with the *fatty acid biosynthetic process, signal transduction*, and *feeding behavior* (Table SIII).

**Figure 5.**
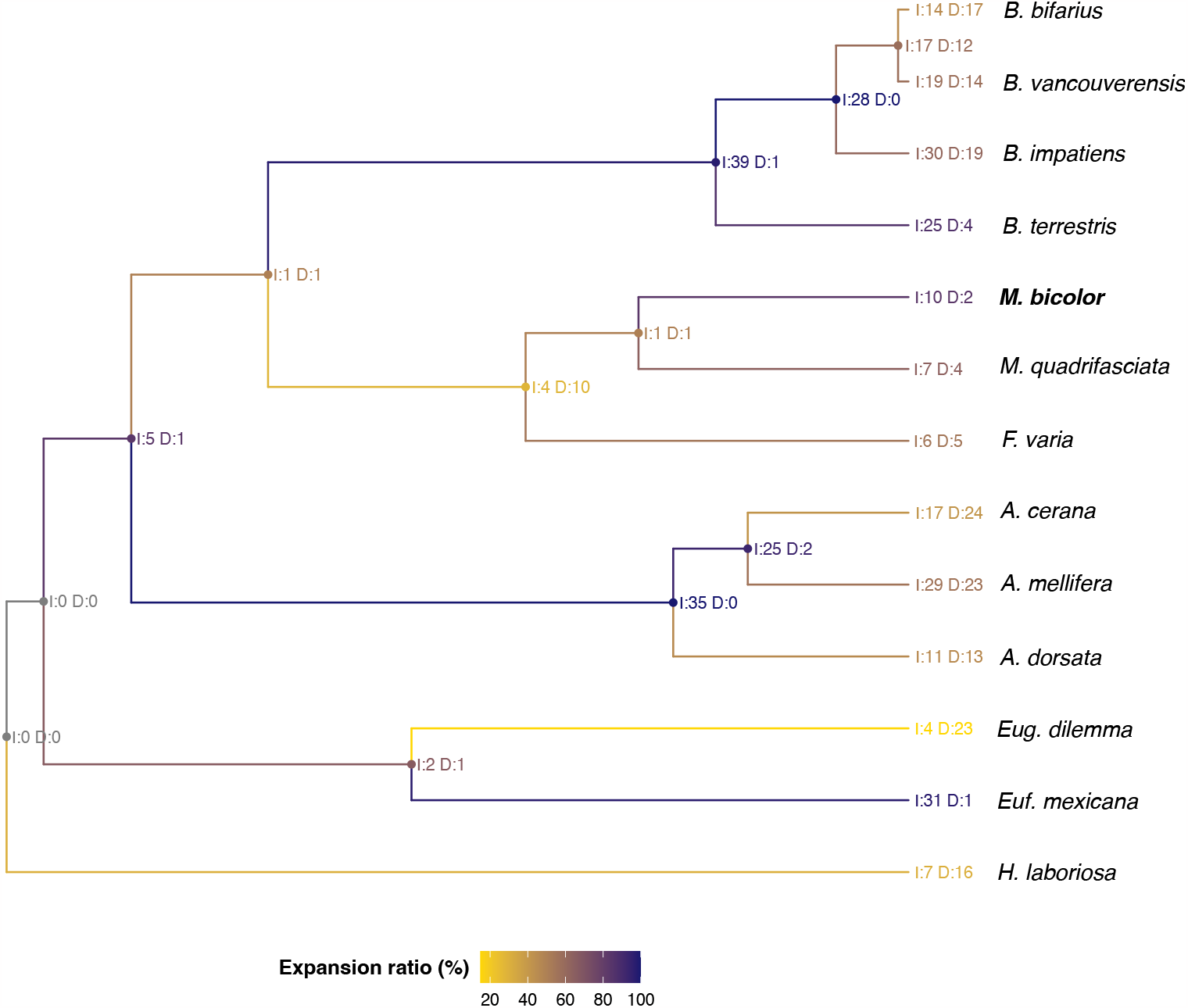
Ultrametric tree of the studied bee species indicating the significant gene family changes in each node. I = gene family expansion events (increase), D = gene family contraction events (decrease). Colors indicate the proportion of changes that were due to gene family expansion, i.e., if the node experienced more gene family expansion events than contractions it will be colored in a colder color as indicated in the color legend. *M. bicolor* is highlighted in bold.

Syntenic analyses among the genomes of *M. bicolor, B. terrestris*, and *A. mellifera* revealed multiple genome rearrangements across this eusocial corbiculate species (Figure 6), with fewer events between *B. terrestris* and *M. bicolor* when compared to *A. mellifera* (Figures S4 and S5, respectively). This is explained by the closer phylogenetic relationship between *B. terrestris* and *M. bicolor* when compared to the honeybee as demonstrated previously through molecular phylogenies [55]. The long scaffold (S1) found in *M. bicolor* is likely correspondent to the larger chromosome 1 [13], as syntenic regions show it has several similarities with the honeybee and the bumblebee chromosome 1. Interestingly, this chromosome also includes syntenic regions in tandem with several chromosomes in the other two species (Figure 6 and S4), suggesting it has gone through multiple rearrangements in Meliponini.

**Figure 6.**
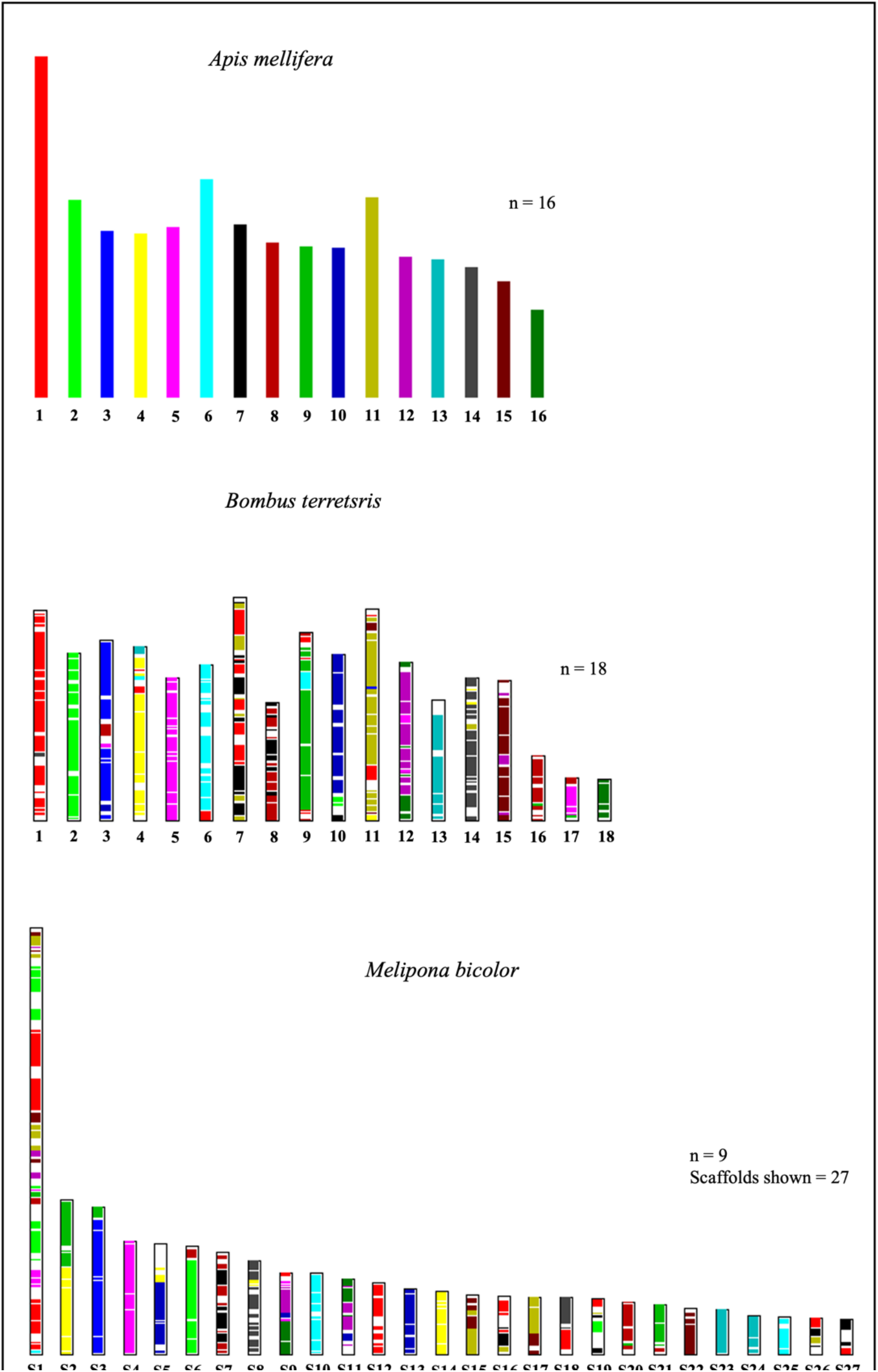
Genomic syntenies inferred among the genome assembly of *A. mellifera* (upper bars), *B. terrestris* (middle bars), and *M. bicolor* (lower bars). The comparisons shown use *A. mellifera* as the color reference, the same colors represent syntenic chromosome regions. For *M. bicolor* the largest 27 scaffolds are presented instead of chromosomes, representing 80% of the total genome assembly.

### Evolution of venom genes in stingless bees

We investigated the molecular evolution of 11 venom protein genes (*venom carboxylesterase-6-like, venom dipeptidyl peptidase 4, venom serine carboxypeptidase, venom peptide isomerase, c1q-like venom protein, cysteine-rich venom protein, venom protease-like, venom allergen* 5, *toxin 3FTx-Lei1, venom serine protease*, and *hyaluronidase*) by estimating synonyms and non-synonym nucleotide changes and patterns of selection (d_N_, d_S,_ and ω values) focused on the Meliponini ancestral node. In the *branch* test, we found all venom genes tested were under negative or purifying selection in all the species studied, including all stingless bees (i.e., **ω** <1 Table SIV). Of these, only two genes, the *venom dipeptidyl peptidase 4* and the *cysteine-rich venom protein*, significantly differed in Meliponini under the *b_free* evolutionary model. The *venom dipeptidyl peptidase 4*, a gene related to toxin maturation [30], had a larger **ω** value than the background data [**ω** (f)=0,302 e **ω** (b)=0,172] indicating a less intense negative selection in this gene. Conversely, the *cysteine-rich venom protein* showed the opposite pattern [**ω** (f)=0,018 e **ω** (b)=0,179] suggesting it could be under stronger positive selection within the Meliponini. The remaining nine venom genes tested were not significantly divergent in the Meliponini. Under the *branch-site* test, only the gene *venom dipeptidyl peptidase 4* was identified with a diverging evolutionary pattern in Meliponini, which suggests codon purifying selection is less intense in this gene among stingless bees, in agreement with the *branch* test.

## Discussion

The stingless bees comprise the most diverse group of highly eusocial bees, they have significant economic and ecological relevance in natural regions and show a range of unique ecological and behavioral adaptations. Still, genomic studies in stingless bees are scarce and we are far from understanding and characterizing the molecular diversity of these bees. Herein, we present a high-quality assembly for the genome of *Melipona bicolor*, the first stingless bee genome assembled using long-read sequencing data, allowing us to recover chromosome length scaffolds. Besides representing a valuable reference for future molecular studies, this assembly contributes to improving our understanding of the genomic diversity of eusocial corbiculate bees as orthologous analyses suggest stingless bees’ genes might be under-estimated in databases.

Syntenic analyses among eusocial corbiculate species and *M. bicolor* have shown interesting evolutionary patterns, such as several chromosome rearrangements among these lineages (Figure 6). Based on the syntenic blocks, we found the longest scaffold in the genome of *M. bicolor* (lengthening 36 Mb) corresponds to chromosome 1 of the other bee genomes compared. Cytogenetic studies in *Melipona* have already shown that chromosome 1 is indeed the longest in most species of this genus – including *M. bicolor* [13, 61]. Here we found that scaffold 1 in *M. bicolor* encompasses regions syntenic to several chromosomes in *B. terrestris* and *A. mellifera*, especially to chromosomes one, two, and eleven (Figures S4 and S5). The coverage and mapping quality of long reads to this scaffold is rather good (Figure S1) strongly suggesting these results are not due to an assembly error. Instead, we argue this finding supports the hypothesis that multiple rearrangements involving several chromosomes occurred during the evolution of the *Melipona* genus resulting in its reduced haploid chromosome number [62]. It is unknown though how conserved this chromosome structure is across the genus, once size variation has been observed and heterochromatin distribution is remarkably variable across *Melipona* species [13, 61].

Compared to the other bees studied here, our gene annotation included more genes and over 35% of these genes were not assigned to Orthogroups. This may indicate that either we have identified many false positive genes in our genome assemblage, or the gene set of other bees is underrepresented for Meliponini genes. Similarly, *M. quadrifasciata* also had one of the smaller proportions of genes included in orthogroups (80.1%). Notably, 15,010 genes in *M. bicolor* had a functional annotation, but only 12,777 genes were assigned to orthogroups. This supports the hypothesis that at least some of the genes in *M. bicolor* are real genes simply not reported in the other bee species, as they have received functional annotations. Moreover, through the BlobToolKit analyses, we found that some contigs significantly blasted to non-corbiculate bees from the Megachilidae and the Andrenidae families (Figure S3). This suggests that including these more distant lineages could have increased orthologous identification. We expect that as new high-quality bee genomes are sequenced the number of genes and orthologs found across species will also increase. Consequently, we argue that improving the completeness and reducing fragmentation of reference genomes, especially in stingless bee genomes, will affect the identification of adaptative orthologous. Therefore, our orthologous comparisons should be regarded as preliminary at this stage.

In the eusocial corbiculate genomes (including Apini, Bombini, and *M. bicolor*), we found 19 exclusive ortholog families that did not occur in Euglossini and *H. laboriosa*. These were related to *nucleosome assembly, phospholipid metabolic process, lipid catabolic process, defense response to bacterium*, and *arachidonic acid secretion*. Additionally, through the gene family change analyses, we found that five gene families were significantly expanding in all eusocial corbiculate. These expanding gene families were related to carbohydrate and lipidic metabolism, and to signaling transduction, which included a gene family of odorant receptor genes. In agreement with these findings, adaptations in the metabolic and energetic pathways have also been reported to be involved with eusocial adaptations in multiple studies [63–67]. We also found a few lineage-specific genes that should be further investigated to explore species adaptations, such as polygyny in *M. bicolor*. This includes 178 genes in species-specific orthogroups found only in *M. bicolor* and 12 gene families that significantly varied in size and included genes involved with the *fatty acid biosynthetic process, signal transduction*, and *feeding behavior*.

The maintenance of venom genes in stingless bee genomes is an intriguing evolutionary question. Although the melittin has been lost in this group, as observed by Koludarov et al. [30] and here, other 11 protein-related genes are still present in their genomes even though stingless bees (as the name suggests) do not have the morphological apparatus necessary for venom injection [9]. In addition, we found that most of these venom genes are still under purifying selection in stingless bees. Only two of the tested genes have significantly diverging molecular evolutionary patterns in stingless bees, and only one of them, the *venom dipeptidyl peptidase 4* is – as it would be expected – evolving through less intense purifying selection in stingless bees when compared to the other stinging species. Unexpectedly, the *cysteine-rich venom protein* showed an even stronger purifying selection in *Meliponini*. The *venom dipeptidyl peptidase 4* is related to toxin maturation [30]. In snakes, the *cysteine-rich venom protein* is involved with the inhibition of smooth muscle contraction and cyclic nucleotide-gated ion channels [68], but its function in bee venom is unknown. Koludarov et al. [30] suggested that most bee venoms may have been co-opted from other physiological functions and that in stingless bees, most of these genes are still present because they would be biologically relevant for other functions, even though they might have lost their venomous role. These findings are corroborated by our analyses, as we found that most venom genes are still evolving under purifying selection in stingless bees, supporting the hypothesis that they would be involved with alternative non-venom-related functions.

## Conclusions

Herein we provide the complete nuclear genome of the stingless bee *Melipona bicolor* that along with the complete mitochondrial genome previously published in (Araujo & Arias, 2019) completes the genomic description of *M. bicolor*, the only true polygynyc bee recognized so far. To illustrate the relevance of this dataset, we performed comparative analyses among the corbiculate bees unrevealing broad patterns of genomic evolution across this clade, including chromosomal rearrangements, gene family expansions, and lineage-specific genes. Finally, we found that the conserved repertoire of venom component genes remains under purifying selection in stingless bees despite their inability to sting. These findings contribute to our understanding of the molecular diversity and adaptation within stingless bees, and as the first high-quality genome available for this group, we believe this data will represent a useful source for future studies.

## Methods

### Data sequencing and genome assembly

We sampled *Melipona bicolor* from colonies kept at the bee lab at the University of São Paulo, São Paulo Campus – Brazil. For the genome sequencing, we used two haploid male pupae that were collected by opening reproductive cells within the colony. The males were recognized due to their characteristic head and eye shape. One male pupa was sequenced in the Illumina platform for paired reads of 100 bp using the TruSeq DNA PCR-Free Fit Library preparation kit. The second male was sequenced using the Pacbio Sequel SMRT Cell 1M and the Sequencing Binding Kit 1.0 to generate long reads. Both sequencing strategies and library preparations were performed by Macrogen at their South Korea facility. For annotation purposes, we additionally sequenced the transcriptome of three females aged between 12-14 days. Upon their emergence, females were color-marked and reintroduced into the colony. After a period of 12-14 days, we retrieved the marked females from the colony and immediately froze them in liquid nitrogen. During this stage of development, female workers are expected to primarily engage in tasks within the colony and are commonly referred to as nurses [31]. Total RNA from three female whole bodies was pooled for RNASeq at the Illumina HiSeq 2500 platform to generate 100bp paired reads. The RNASeq was performed at Lactad - Unicamp. Then, the *de novo* transcriptome assembly was performed using Trinity v2.8.4 [32] to be used in the genome annotation. The genome assembly was initially performed in two ways, first using Falcon v1.3.0 [33] with only the long-read data, and secondly using MaSuRCA v3.3.5 [34] based on a combination of short and long reads. Parameters used in the Falcon assembly were: *genome_size=0; seed_coverage=20; length_cutoff=1000; ovlp_daligner_option=--e*.*93 -l2500 -k24 -h1024 -w6 -s100*. MaSuRCA run was set up for a k-mer size of 67 for the graph step and 25x coverage for the longest reads. Both assemblies were corrected using arrow (Pacbio tools gcpp v1.9.0) with default parameters after realignment of the long reads with pbmm2 v1.2.0. After polishing the two assemblies were combined using quickmerge wrapper v0.3 with MUMmer v3.23 [35] with the parameter *-l 721765*. We then removed the mitochondrial contig from the assembly by aligning *M. bicolor* mitogenome[12] to the assembly using Last v1047 [36] with the options *-uNEAR -R01*. For further scaffolding, we used the transcriptome assembly and the program SCUBAT v2 [37] with 40,000 bp as the maximum intron size, since 99% of all introns assembled had a maximum size of 38,984 bp. The resulting assembly was polished with arrow again, using only the long reads and default parameters, then with pilon v1.23 [38] using the parameter *-Xmx250G* and the short reads. We then re-aligned the long and short reads using pbmm2 and bwa v0.7.17 [39], respectively, to the polished assembly and analyzed the quality of the alignments using Qualimap v2.2.1 [40]. Based on this, we removed contigs with extreme coverage bias (i.e., with mean coverage ≤ 1) according to long reads alignment, and one contig that had a very high coverage and aligned to the mitogenome (GC content 40%). The removal of these contigs did not affect BUSCO quality scores. Lastly, an additional polishing step using short reads and long reads combined was performed using hypo v1.0.3 [41]. For quality estimation throughout the assembly processes, we have used QUAST v5.0.2 [42], BUSCO v5.1.2 [43], Qualimap, and BlobToolKit v4.1.5 [44].

### Genome annotation

Repetitive elements were identified in the genome using the pipeline available at https://github.com/nat2bee/repetitive_elements_pipeline. Briefly, we used RepeatModeler v1.0.11 [45], TransposonPSI [46], and LTRharvest from GenomeTools v1.6.1 [47] to build custom repeat libraries. These libraries were merged into a single non-redundant library of repetitive elements for *M. bicolor* using USEARCH v11.0.667 [48] (<80% identity). RepeatClassifier was used for the library annotation. Then, we concatenated our custom library with the Dfam v3.1 Hymenoptera library included in RepeatMasker v4.1.0 [49] and used the same program to annotate and soft mask the repeats found in the genome based on our custom library (available at https://github.com/nat2bee/repetitive_elements_pipeline). Repeats Summary statistics of the annotated repeats were obtained using a custom script *RepeatMasker_stats*.*py* (https://github.com/nat2bee/repetitive_elements_pipeline). Genome annotation of non-masked regions was performed using Funannotate v1.7.4 pipeline [50] which combined gene predictions based on *M. bicolor de novo* transcriptomic assembly (9,458 gene models) and the Insecta gene model database from BUSCO (all database versions are detailed in Supplementary File 1). Functional annotation of the genes was performed also under Funannotate in tandem with InterProScan5. *M. bicolor* genes were additionally blasted, using NCBI-blatsp [51], against bee proteins in the UniRef90 database (from August 2021). From these results gene names were preferentially assigned and the gene ontology terms (GO) were retrieved. Finally, the annotation and the genome were trimmed according to NCBI’s quality filtering.

### Comparative genomics

We compared the genome assembly of *M. bicolor* with that of other eleven corbiculate bees (*Apis cerana, A. dorsata, A. mellifera, Bombus bifarius, B. impatiens, B. terrestris, B. vancouverensis, Eufriesea mexicana, Euglossa dilemma, Frieseomelitta varia*, and *Melipona quadrifasciata*) and one outgroup species (*Habropoda laboriosa*) using OrthoFinder v2.5.4 [52] and CAFE v5.0.0 [53]. These genomes were chosen because they comprise lineages representing all corbiculate clades and were available in the NCBI database along with their attributed gene annotations (Table S1), except for *E. dilemma* whose genome annotation was retrieved from the Beebase (https://hymenoptera.elsiklab.missouri.edu/beebase/consortium_datasets). For the genomes in which the RefSeq annotation was not available (i.e., for *E. dilemma, F. varia, M. quadrifasciata*, and *M. bicolor*), we selected one isoform per gene by grouping similar proteins using CD-Hit V4.8.1 [54] with a 70% similarity threshold. Using OrthoFinder with default parameters we generated the species tree and re-rooted it at *H. laboriosa* to identify orthologs and species-specific genes across all branches. We then generated an ultrametric tree using the OrthoFinder *make_ultrametric*.*py* script and calibrated the tree root node at 105 million years [55]. We identified gene family changes across the tree nodes using CAFE. We initially removed the two largest gene families using the *lade_and_size_filter*.*py* script and then estimated the parameters to run CAFE. First by measuring the error parameter to account for possible genome assembly errors, then by testing multiple k values (from 1 to 10) to choose the one with the highest likelihood score, and finally, by running the estimates multiple times and getting the median lambda values. We finally run CAFE using the command *-eerror_model_017*.*txt -l 0*.*0017312* where the error model represents a text file containing the following information: *maxcnt: 105; cntdiff: -1 0 1; 0 0 0*.*982961 0*.*0170389; 1 0*.*0170389 0*.*965922 0*.*0170389*.

Since *M. bicolor* assembly resulted in large contigs and N50, we inferred the synteny between this genome and two high-quality genome assemblies representing the other eusocial corbiculate bee lineages – *i*.*e*., the Apini (*A. mellifera*) and Bombini (*B. terrestris*) – both assemblies with associated chromosome mappings. For this analysis, we first blasted species protein sets with each other using blastp from NCBI-blast v2.9.0 [51] with the parameters *-e 1e-10 -b 5 -v 5 -m 8*, and then used the MCScanX [56] with default parameters to identify and illustrate collinear blocks across species. Since the genome of *M. bicolor* is not assembled to the chromosome level, we inferred the synteny based on the 27 largest scaffolds, which corresponds to 80% of the genome.

### Evolution of venom genes in stingless bees

To study the evolution of venom-related genes in stingless bees, we selected the venom-associated genes based on Koludarov et al. (2022) findings. These authors focused on 12 proteins prevalent in the bee venom. Among these, we excluded the *melittin* gene from our analysis because it is not present in the stingless bee genomes (including *M. bicolor*). Orthogroups containing the remaining 11 venom protein genes (i.e., *venom carboxylesterase-6-like, venom dipeptidyl peptidase 4, venom serine carboxypeptidase, venom peptide isomerase, c1q-like venom protein, cysteine-rich venom protein, venom protease-like, venom allergen* 5, *toxin 3FTx-Lei1, venom serine protease*, and *hyaluronidase*) were filtered from OrthoFinder results obtained in the comparative analyses. To select only a single gene copy per orthogroup we aligned all sequences included in the orthogroup using Mafft v7.310 [57] – with the parameter *adjustdirectionaccurately* – and selected the best-aligning gene copy (*i*.*e*., the copy whose alignment produced fewer gaps in comparison to the other bee species).

The selected amino acid sequences from all species were aligned with Mafft and posteriorly polished using GBlocks v0.91b [58]. Molecular evolution tests supported by species phylogeny were performed using CODEML within the PAML4 package [59] to estimate synonyms and non-synonym nucleotide changes (d_N_, d_S,_ and ω values). ETE3 v3.1.2 [60] was used to automate these analyses. The Meliponini ancestral node was used as the foreground node to focus on the evolutionary pattern of venom genes in this branch. We tested six different evolutionary models for each gene, three using the *branch* (*M0, b_neut*, and *b_free*) and three using the *branch-site* (*M1, bsA1* e *bsA*) model. The *branch* model tests estimate the molecular evolution of the entire gene in the foreground node compared to the remaining (background) nodes, while the branch-site models search for codon positive selection in the foreground node [59]. Model results were interpreted according to likelihood ratio tests and associated p-values, which were corrected for multiple comparisons with a cut-off of p < 0.05.

## Notes

https://github.com/nat2bee/repetitive_elements_pipeline

### Competing Interest Statement

The authors have declared no competing interest.

## References

1. Jaffé R, Pope N, Acosta AL, Alves DA, Arias MC, D. la Rúa P, et al. Beekeeping practices and geographic distance, not land use, drive gene flow across tropical bees. Mol Ecol. 2016;25:5345–58.

2. Garibaldi LA, Carvalheiro LG, Vaissiere BE, Gemmill-Herren B, Hipolito J, Freitas BM, et al. Mutually beneficial pollinator diversity and crop yield outcomes in small and large farms. Science (1979). 2016;351:388– 91.

3. Bueno FGB, Kendall L, Alves DA, Tamara ML, Heard T, Latty T, et al. Stingless bee floral visitation in the global tropics and subtropics. Glob Ecol Conserv. 2023;43:e02454.

4. Lopes AV, Porto RG, Cruz-Neto O, Peres CA, Viana BF, Giannini TC, et al. Neglected diversity of crop pollinators: Lessons from the world’s largest tropical country. Perspect Ecol Conserv. 2021;19:500–4.

5. Potts SG, Imperatriz-Fonseca V, Ngo HT, Aizen MA, Biesmeijer JC, Breeze TD, et al. Safeguarding pollinators and their values to human well-being. Nature. 2016;540:220–9.

6. Dicks L V., Viana B, Bommarco R, Brosi B, Arizmendi M del C, Cunningham SA, et al. Ten policies for pollinators. Science (1979). 2016;354:975–6.

7. Alves DA, George EA, Kaur R, Brockmann A, Hrncir M, Grüter C. Diverse communication strategies in bees as a window into adaptations to an unpredictable world. Proc Natl Acad Sci U S A. 2023;120:e2219031120.

8. Freitas FC de P, Lourenço AP, Nunes FMF, Paschoal AR, Abreu FCP, Barbin FO, et al. The nuclear and mitochondrial genomes of Frieseomelitta varia - a highly eusocial stingless bee (Meliponini) with a permanently sterile worker caste. BMC Genomics. 2020;21:386.

9. Michener CD. The bees of the world. second. Johns Hopkins University Press; 2007.

10. Silveira FA, Melo GAR, Almeida EAB. Abelhas brasileiras: Sistemática e identificação. First edit. Belo Horizonte; 2002.

11. NCBI genome search. https://www.ncbi.nlm.nih.gov/genome/?term=meliponini. 2023.

12. Araujo NS, Arias MC. Mitochondrial genome characterization of Melipona bicolor: Insights from the control region and gene expression data. Gene. 2019;705 April:55–9.

13. Rocha MP, Pompolo S das G, Dergam JA, Fernandes A, Campos LA de O. DNA characterization and karyotypic evolution in the bee genus Melipona (Hymenoptera, Meliponini). Hereditas. 2002;136:19–27.

14. Tavares MG, Carvalho CR, Soares FAF. Genome size variation in Melipona species (Hymenoptera: Apidae) and sub-grouping by their DNA content. Apidologie. 2010;41:636–42.

15. Velthuis HHW, Roeling A, Imperatriz-Fonseca VL. Repartition of reproduction among queens in the polygynous stingless bee Melipona bicolor. In: Proceedings of the Section Experimental and Applied Entomology of the Netherlands Entomological Society (NEV). 2001. p. 45–9.

16. Velthuis HHW, De Vries H, Imperatriz-Fonseca VL. The polygyny of Melipona bicolor: Scramble competition among queens. Apidologie. 2006;37:222–39.

17. Reis EP dos, Campos LA de O, Tavares MG. Prediction of social structure and genetic relatedness in colonies of the facultative polygynous stingless bee Melipona bicolor (Hymenoptera, Apidae). Genetics and Molecular Biology Online. 2011.

18. Cepeda OI. Division of labor during brood production in stingless bees with special reference to individual participation. Apidologie. 2006;37:175–90.

19. Velthuis HHW, Vries H de, Imperatriz-Fonseca VL. The Polygyny of Melipona bicolor: Scramble Competition Among Queens. Apidologie. 2006;37:222–39.

20. Michener CD. The Bees of the World. Baltimore and London: The Johns Hopkins University Press; 2000.

21. Casale TB, Burks AW. Hymenoptera-Sting Hypersensitivity. New England Journal of Medicine. 2014;370:1432–9.

22. de Graaf DC, Aerts M, Danneels E, Devreese B. Bee, wasp and ant venomics pave the way for a component-resolved diagnosis of sting allergy. J Proteomics. 2009;72:145–54.

23. Kim W. Bee Venom and Its Sub-Components: Characterization, Pharmacology, and Therapeutics. Toxins (Basel). 2021;13.

24. Kim BY, Kim YH, Park MJ, Yoon HJ, Lee KY, Kim HK, et al. Dual function of a bumblebee (Bombus ignitus) serine protease inhibitor that acts as a microbicidal peptide and anti-fibrinolytic venom toxin. Dev Comp Immunol. 2022;135:104478.

25. El Mehdi I, Falcão SI, Boujraf S, Mustapha H, Campos MG, Vilas-Boas M. Analytical methods for honeybee venom characterization. J Adv Pharm Technol Res. 2022;13:154–60.

26. Barkan NP, Chevalier M, Pradervand JN, Guisan A. Alteration of Bumblebee Venom Composition toward Higher Elevation. Toxins (Basel). 2019;12.

27. Pucca MB, Cerni FA, Oliveira IS, Jenkins TP, Argemí L, Sørensen C V., et al. Bee Updated: Current Knowledge on Bee Venom and Bee Envenoming Therapy. Front Immunol. 2019;10.

28. Duda TF, Chang D, Lewis BD, Lee T. Geographic Variation in Venom Allelic Composition and Diets of the Widespread Predatory Marine Gastropod Conus ebraeus. PLoS One. 2009;4:e6245.

29. Menezes MC, Furtado MF, Travaglia-Cardoso SR, Camargo ACM, Serrano SMT. Sex-based individual variation of snake venom proteome among eighteen Bothrops jararaca siblings. Toxicon. 2006;47:304–12.

30. Koludarov I, Velasque M, Timm T, Greve C, Ben Hamadou A, Kumar Gupta D, et al. A common venomous ancestor?Prevalent bee venom genes evolved before the aculeate stinger while few major toxins are bee-specific. bioRxiv. 2022. 10.1101/2022.01.21.477203.

31. Mateus S, Ferreira-Caliman MJ, Menezes C, Grüter C. Beyond temporal-polyethism: division of labor in the eusocial bee Melipona marginata. Insectes Soc. 2019;66:317–28.

32. Grabherr MG, Haas BJ, Yassour M, Levin JZ, Thompson DA, Amit I, et al. Trinity: reconstructing a fulllength transcriptome without a genome from RNA-Seq data. Nat Biotechnol. 2013;29:644–52.

33. Chin C-S, Peluso P, Sedlazeck FJ, Nattestad M, Concepcion GT, Clum A, et al. Phased diploid genome assembly with single-molecule real-time sequencing. Nat Methods. 2016;13:1050–4.

34. Zimin A V., Marçais G, Puiu D, Roberts M, Salzberg SL, Yorke JA. The MaSuRCA genome assembler. Bioinformatics. 2013;29:2669–77.

35. Chakraborty M, Baldwin-Brown JG, Long AD, Emerson JJ. Contiguous and accurate de novo assembly of metazoan genomes with modest long read coverage. Nucleic Acids Res. 2016;44:e147.

36. Kiełbasa SM, Wan R, Sato K, Horton P, Frith MC. Adaptive seeds tame genomic sequence comparison. Genome Res. 2011;21:487–93.

37. Koutsovoulos G, Kao D. SCUBAT2: Scaffolding Contigs Using BLAST and Transcripts v2. 2016.

38. Walker BJ, Abeel T, Shea T, Priest M, Abouelliel A, Sakthikumar S, et al. Pilon: An Integrated Tool for Comprehensive Microbial Variant Detection and Genome Assembly Improvement. PLoS One. 2014;9:e112963.

39. Li H. Aligning sequence reads, clone sequences and assembly contigs with BWA-MEM. 2013.

40. García-Alcalde F, Okonechnikov K, Carbonell J, Cruz LM, Götz S, Tarazona S, et al. Qualimap: evaluating next-generation sequencing alignment data. Bioinformatics. 2012;28:2678–9.

41. Kundu R, Casey J, Sung W-K. HyPo: Super Fast & Accurate Polisher for Long Read Genome Assemblies. bioRxiv. 2019;:2019.12.19.882506.

42. Gurevich A, Saveliev V, Vyahhi N, Tesler G. QUAST: quality assessment tool for genome assemblies. Bioinformatics. 2013;29:1072–5.

43. Simão FA, Waterhouse RM, Ioannidis P, Kriventseva E V., Zdobnov EM. BUSCO: Assessing genome assembly and annotation completeness with single-copy orthologs. Bioinformatics. 2015;31:3210–2.

44. Challis R, Richards E, Rajan J, Cochrane G, Blaxter M. BlobToolKit – Interactive Quality Assessment of Genome Assemblies. G3: Genes|Genomes|Genetics. 2020;10:1361.

45. Smit A, Hubley R. RepeatModeler Open-1.0. 2010.

46. Haas B. TransposonPSI: An Application of PSI-Blast to Mine (Retro-)Transposon ORF Homologies. 2010.

47. Ellinghaus D, Kurtz S, Willhoeft U. LTRharvest, an efficient and flexible software for de novo detection of LTR retrotransposons. BMC Bioinformatics. 2008;9.

48. Edgar RC. Search and clustering orders of magnitude faster than BLAST. Bioinformatics. 2010;26:2460–1.

49. Smit A, Hubley R, Green P. RepeatMasker Open. 2013.

50. Palmer J, Stajich J. Funannotate. 2019.

51. Camacho C, Coulouris G, Avagyan V, Ma N, Papadopoulos J, Bealer K, et al. BLAST+: architecture and applications. BMC Bioinformatics. 2009;10:421.

52. Emms DM, Kelly S. OrthoFinder: Phylogenetic orthology inference for comparative genomics. Genome Biol. 2019;20:238.

53. De Bie T, Cristianini N, Demuth JP, Hahn MW. CAFE: a computational tool for the study of gene family evolution. Bioinformatics. 2006;22:1269–71.

54. Huang Y, Niu B, Gao Y, Fu L, Li W. CD-HIT Suite: A web server for clustering and comparing biological sequences. Bioinformatics. 2010;26:680–2.

55. Bossert S, Murray EA, Almeida EAB, Brady SG, Blaimer BB, Danforth BN. Combining transcriptomes and ultraconserved elements to illuminate the phylogeny of Apidae. Mol Phylogenet Evol. 2019;130 July 2018:121– 31.

56. Wang Y, Tang H, Debarry JD, Tan X, Li J, Wang X, et al. MCScanX: a toolkit for detection and evolutionary analysis of gene synteny and collinearity. Nucleic Acids Res. 2012;40:e49.

57. Katoh K, Standley DM. MAFFT Multiple Sequence Alignment Software Version 7: Improvements in Performance and Usability. Mol Biol Evol. 2013;30:772–80.

58. Talavera G, Castresana J. Improvement of Phylogenies after Removing Divergent and Ambiguously Aligned Blocks from Protein Sequence Alignments. Syst Biol. 2007;56:564–77.

59. Yang Z. PAML 4: Phylogenetic Analysis by Maximum Likelihood. Mol Biol Evol. 2007;24:1586–91.

60. Huerta-Cepas J, Serra F, Bork P. ETE 3: Reconstruction, Analysis, and Visualization of Phylogenomic Data. Mol Biol Evol. 2016;33:1635–8.

61. da Cunha MS, Travenzoli NM, Ferreira R de P, Cassinela EK, da Silva HB, Oliveira FPM, et al. Comparative cytogenetics in three Melipona species (Hymenoptera: Apidae) with two divergent heterochromatic patterns. Genet Mol Biol. 2018;41:806–13.

62. Travenzoli NM, Cardoso DC, De Azevedo Werneck H, Fernandes-Salomão TM, Tavares MG, Lopes DM. The evolution of haploid chromosome numbers in Meliponini. PLoS One. 2019;14.

63. Araujo N de S, Arias MC. Gene expression and epigenetics reveal species-specific mechanisms acting upon common molecular pathways in the evolution of task division in bees. Sci Rep. 2021;11:3654.

64. Chandra V, Fetter-Pruneda I, Oxley PR, Ritger AL, McKenzie SK, Libbrecht R, et al. Social regulation of insulin signaling and the evolution of eusociality in ants. Science (1979). 2018;361:398–402.

65. Fischer EK, O’Connell LA. Modification of feeding circuits in the evolution of social behavior. J Exp Biol. 2017;220:92–102.

66. Morandin C, Tin MMY, Abril S, Gómez C, Pontieri L, Schiøtt M, et al. Comparative transcriptomics reveals the conserved building blocks involved in parallel evolution of diverse phenotypic traits in ants. Genome Biol. 2016;17:1–19.

67. Weitekamp CA, Libbrecht R, Keller L. Genetics and Evolution of Social Behavior in Insects. Annu Rev Genet. 2017;51:219–39.

68. Yamazaki Y, Morita T. Structure and function of snake venom cysteine-rich secretory proteins. Toxicon. 2004;44:227–31.

